# Monosynaptic ventral tegmental area glutamate projections to the locus coeruleus enhance aversive processing

**DOI:** 10.1101/2024.10.04.615025

**Authors:** Kyle E. Parker, Chao-Cheng Kuo, Alex R. Buckley, Abigail P. Patterson, Vincent Duong, Sarah C. Hunter, Jordan G. McCall

## Abstract

Distinct excitatory synaptic inputs to the locus coeruleus (LC) modulate behavioral flexibility. Here we identify a novel monosynaptic glutamatergic input to the LC from the ventral tegmental area (VTA). We show robust VTA axonal projections provide direct glutamatergic transmission to LC. Despite weak synaptic summation, optogenetic activation of these axons enhances LC tonic firing and facilitates real-time and conditioned aversive behaviors. We hypothesized this projection may modulate synaptic integration with other excitatory inputs. We then used coincident VTA-LC photostimulation with local electrical stimulation and observed enhanced LC burst induction. To determine whether this integration also occurs *in vivo*, we took an analogous approach measuring reward-seeking behavior during unpredictable probabilistic punishment. Here, glutamatergic VTA-LC photostimulation during a concurrent noxious stimulus did not delay reward-seeking behavior, but increased probability of task failure. Together, we identified a novel VTA-LC glutamatergic projection that drives concurrent synaptic summation during salient stimuli to promote behavioral avoidance.

## Introduction

Excitation of the locus coeruleus-noradrenergic (LC-NE) system greatly influences many distinct behaviors, including arousal, learning, nociception, stress, and negative affect. Glutamatergic excitation of LC-NE neurons, in particular, is often associated with behavioral flexibility and attention shifting to salient stimuli^1–3^. However, glutamate signaling in the LC-NE system has also been implicated in modulating negative affective states including neuropathic pain^4,5^, opioid-withdrawal^6–10^, depression^11^, and anxiety^12,13^. Furthermore, multiple excitatory and modulatory inputs to LC-NE neurons have been shown to enhance LC-NE activity in several LC-mediated negative affective states^8,14–22^. Despite rigorous study of the downstream changes in NE release^10,19,23–27^, relatively little is known about integration of coincident glutamatergic afferents on LC-NE neurons and their potential negative affective consequences. Here we identify a largely unstudied afferent input to the LC from glutamatergic neurons in the ventral tegmental area (VTA), a critical region for reward processing and motivation. Given the molecular, anatomical, and functional heterogeneity of VTA neurons^28–51^ and the role that glutamatergic VTA neurons have in reward and aversive conditioning^36–39,41–49,52,53^, it is imperative to understand the role of this distinct projection among the complex architecture of the VTA efferent system. While projections from glutamatergic VTA neurons to the nucleus accumbens and lateral habenula have opposing valences that can bidirectionally modulate reward and aversion^37,45^, we hypothesized that activation of VTA glutamate projections to the LC drive aversive behaviors. To test this hypothesis, we used *ex vivo* electrophysiology and *in vivo* optogenetics to demonstrate that LC-NE neurons become more responsive to aversive stimuli through summation of glutamatergic transmission from the VTA with coincident excitatory inputs to enhance natural avoidance behaviors that guide actions away from risk-associated outcomes.

## Results

### Monosynaptic glutamatergic transmission from the VTA to LC

Cell type-selective approaches have greatly enhanced our understanding of the molecular, anatomical, and functional heterogeneity of VTA neurons^28–50^. In particular, VTA glutamatergic neurons have clear roles in reward and aversive conditioning^36–39,41–49^. Here, complex VTA projection architecture dictates function, such that modulation of glutamatergic VTA cell bodies is positively reinforcing through activation of nucleus accumbens-projecting VTA-dopamine neurons^38^, but that direct projections to the accumbens or lateral habenula produce aversion^37,45^. While afferent input from the VTA to the LC was first described more than 40 years ago^54^, the projection itself has not been thoroughly interrogated – a striking divergence from the study of either the VTA or LC, both of which have been independently heavily dissected in the intervening years. Early studies suggested a minority of these projections were dopaminergic^55^, but other reports indicated heavier dopamine input^56,57^. The first functional study of this projection showed stimulation of VTA-LC neurons likely recruit LC activity and promote NE release in the prefrontal cortex^58^. This excitation, however, was presumed to occur via dopamine, but neither dopamine nor glutamate receptor antagonists were tested. To determine whether VTA glutamate neurons project directly to the LC, we used a combination of immunohistochemistry and *ex vivo* electrophysiology. First, we selectively expressed YFP-tagged channelrhodopsin-2 (ChR2-eYFP) in glutamatergic VTA neurons, by injecting AAV5-EF1α-DIO-ChR2-eYFP unilaterally into the VTA of *vglut2*^Cre^ mice (*vglut2*^VTA-LC^) (**Fig.1A**). Nine weeks later, ChR2-eYFP is clearly expressed in the VTA with visible bilateral innervation of LC cell bodies and dendrites. This bilateral innervation appears to more robustly target the LC ipsilateral to the VTA injection (**Fig. 1B&C**). To further dissect the excitatory neural population within the VTA, we examined the distribution of CaMKIIα-expressing neurons in VTA following injection of both AAV5-EF1α-DIO-eYFP and AAV5-CaMKIIα-mCherry into the VTA of *vglut2*^Cre^ mice (**Fig. S1A**). In this case, most TH-immunoreactive neurons in VTA also express CaMKIIα (mCherry) and around 65% of them are glutamatergic as indicated by co-expression of Cre-dependent eYFP (**Fig. S1B-D**). To determine whether the collocated VTA fibers make functional synaptic connections with LC neurons, we performed *ex vivo* whole-cell recordings of LC-NE neurons following the criteria described in previous observations^59^. The LC was identified as a transparent region underneath the lateral border of 4^th^ ventricle and LC-NE spindle-shaped neurons were visually identified with size around 25μm (**Fig. 1D-F**). Hyperpolarized current injections caused delayed restoration of spontaneous firing of LC-NE neurons in I-Clamp recording after hyperpolarized current injection **(Fig. S2**). Optogenetic photostimulation (470 nm light pulses with 2 ms duration at 10mW/mm^2^) of ChR2-expressing *vglut2*^VTA-LC^ terminals evoked EPSCs (oEPSC) in LC neurons (**Fig. 1G)**. These oEPSCs appear to be driven by direct monosynaptic inputs as the currents remained intact in the presence of blockers for voltage-gated channels passing Na^+^ and K^+^ ions, to restrict AP generation to axonal components (1μM tetrodotoxin (TTX) and 200μM 4-Aminopyridine). TTX-resistant oEPSCs were completely abolished by antagonists targeting AMPA and NMDA receptors (20 μM DNQX and 50 μM APV, respectively; **Fig. 1G&H**). No IPSCs were observed while membrane potential was held at −40mV under a full blockade of ionotropic glutamatergic receptors suggesting a pure monosynaptic glutamatergic transmission from VTA to LC. Interestingly, TTX and 4AP together increased synaptic latency (**Fig. 1I**) and potentiated oEPSC amplitude (**Fig. 1J)**. These observations may be due to expression of A type K^+^ channels on post-synaptic LC neurons and changes in vesicle release dynamics^60,61^. Consistent with our anatomical results, oEPSCs were observed in the majority of ipsilateral LC-NE neurons (18 out of 20 cells) and roughly half of recorded contralateral neurons (10 out of 18 cells) (**Fig. 1K**). Paired-pulse stimulation (50 ms interstimulus interval) revealed larger amplitude and lower paired-pulse ratios in the ipsilateral side compared to contralateral recordings (**Fig. 1L-N**), suggesting possibly lower release probability, but this could also be explained by possibly less ChR2 expression due to further trafficking in the contralateral projection. It should also be noted that photostimulation-evoked paired-pulse ratios can be underestimates due to the additional calcium influx through axonal ChR2^62,63^. Somewhat surprisingly, repeated photostimulation (10 pulses at 20 Hz) failed to drive transient burst activity due to limited summation from evoked oEPSCs and cleared multi-pulse depression (**Fig. 1O&P**). Together, these results corroborate a distinct, monosynaptic glutamatergic VTA innervation of the LC.

**Figure 1.**
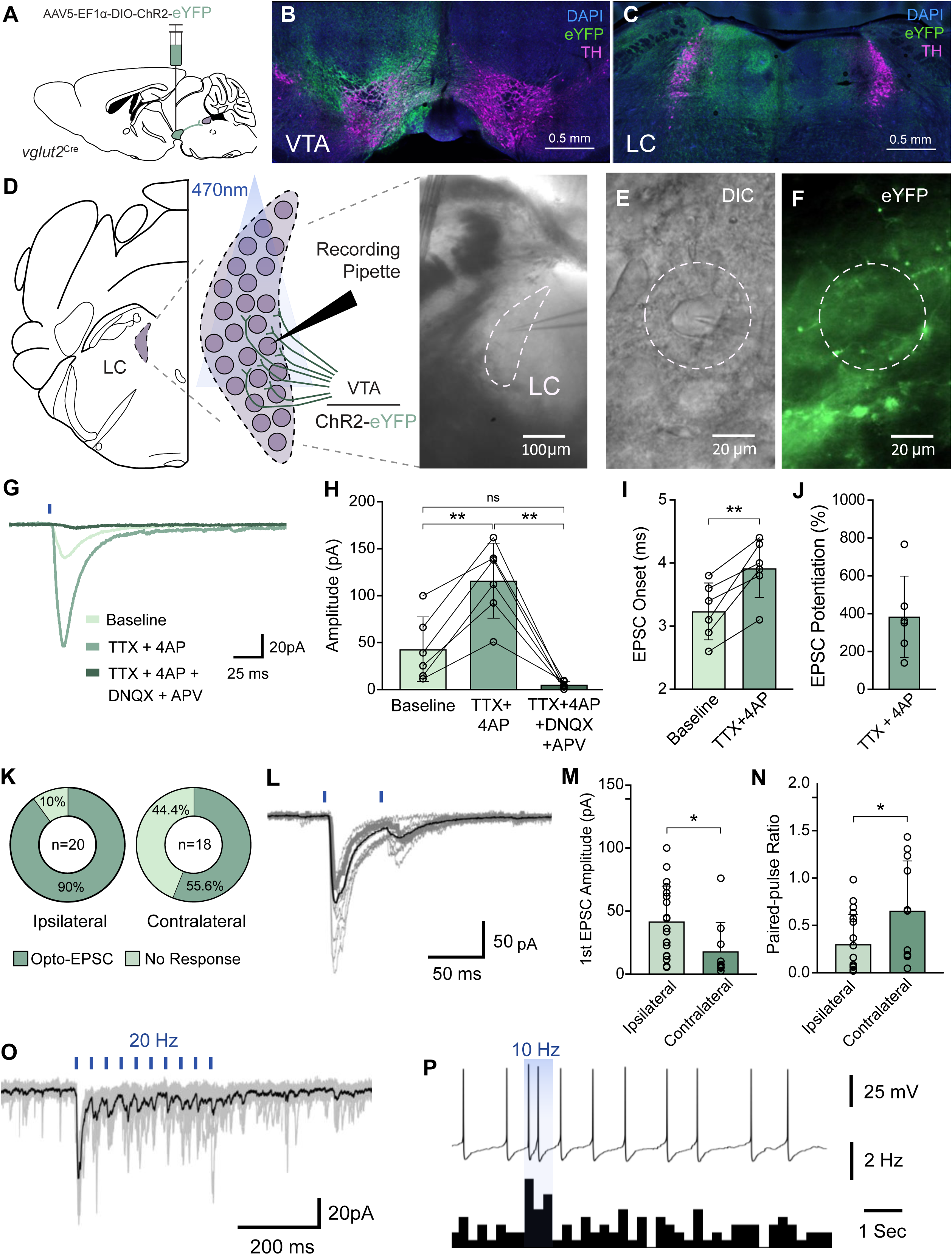
VTA glutamate neurons make monosynaptic inputs onto LC neurons. (**A**) Schematic depicting viral injection of ChR2-eYFP into the VTA of *vglut2*^Cre^ mice. (**B&C**) Representative fluorescent images showing TH- and eYFP-immunoreactive components in VTA (B) and the ipsilateral and contralateral LC (C) (eYFP: green, DAPI: blue, TH: magenta). (**D**) Schematic illustrating the electrophysiological recordings (left) and representative live DIC image of LC (right). (**E**&**F**) DIC and fluorescent images showing a recorded LC-NE neuron surrounded by eYFP-expressing axons. (**G**) Representative traces from pharmacological application during photostimulation-induced glutamatergic transmission onto LC-NE cells. (**H**) Application of TTX + 4AP amplified photostimulation-induced oEPSC amplitude that was nearly entirely abolished by DNQX + APV application. Baseline: 42.8±34.3, TTX+4AP: 115.8±39.9, TTX+4AP+DNQX+APV: 5.2±3.5 pA (Repeated measures One-way ANOVA, F (1.686, 8.432) = 31.50, *p* = 0.0002, Tukey’s multiple comparisons: Baseline vs. TTX + 4AP, p = 0.0035; TTX + 4AP vs. TTX+4AP + DNQX + APV, *p* = 0.0028). (**I**) Application of TTX+4AP significantly increased synaptic latency (3.23±0.45 to 3.92±0.46 ms) and (**J**) potentiated oEPSC amplitude (384±214%) (Paired t test: t(5) = 4.965, *p* = 0.0042). (**K**) Proportion of recorded cells with detectable oEPSCs among ipsilateral (left) and contralateral (right) LC. (**L**) Representative traces of paired-pulse stimulation. (**M&N**) Higher 1^st^ amplitudes in paired-pulse optical illumination in ipsilateral side as well as lower paired-pulse ratio (M, unpaired t test: t(27) = 2.299, *p* = 0.0295; N, Mann-Whitney test: U = 512.299, median (ipsi) = 0.1123, median (contra) = 0.6534, *p* = 0.0311). (**O**) Representative voltage-clamp trace during 10 pulses at 20 Hz. (**P**) Representative current-clamp recording (top) and histogram (bottom) showing increased firing rate during 10 pulses at 10 Hz.

### VTA–LC glutamatergic projection stimulation drives negative affective behaviors

After identifying monosynaptic excitatory input onto LC neurons, we next sought to determine whether this glutamatergic projection drives negative affective behaviors. To do so, we injected AAV5-EF1α-DIO-eYFP or AAV5-EF1α-DIO-ChR2-eYFP into the VTA of *vglut2*^Cre^ mice and implanted a fiber optic above the LC (**Fig. 2A**). We first used an optogenetic real-time place test (RTPT) that initiates photostimulation when the animal enters a stimulation-paired chamber with no other salient environmental stimuli. This enables a quick test to determine whether the stimulation carries a positive or negative behavioral valence (**Fig. 2B**). Here, we found that increasing photostimulation frequency of *vglut2*^VTA-LC^ terminals produced a significant real-time avoidance behavior without affecting total distance moved during exploration (**Fig. 2C-E**). Further examination in C57BL/6J mice injected with AAV5-CaMKIIα-ChR2-eYFP into the VTA and fiber implants in the LC revealed similar behavior to *vglut2*^Cre^ mice (**Fig. S3A&B**). Mice expressing ChR2 in excitatory VTA neurons also showed a frequency-dependent decrease in the percentage of time spent in the stimulation-paired side during RTPT (**Fig. S3B**). Additionally, 20 Hz stimulation reduced the percentage of time spent in the elevated plus maze (EPM) (**Fig. S3C**) a measure of anxiety-related exploratory behavior. Interestingly, the same stimulation did not have any anxiogenic effect in the EPM for *vglut2*^VTA-LC^ mice (**Fig. S3D**), suggesting a functional difference between the CaMKIIα- and *vglut2-*expressing VTA-LC projections. Further, we used the conditioned place preference test to determine whether the previously identified photostimulation-induced negative valence could integrate with environmental stimuli to drive conditioned avoidance behavior. Here, *vglut2*^Cre^ mice again received injection of either AAV5-EF1a-DIO-eYFP or AAV5-EF1a-DIO-ChR2-eYFP into the VTA and a fiber optic implant above the LC (**Fig. 2F**). Mice were allowed to explore all chambers of the apparatus during a pre-test day, then received context-paired 20 Hz photostimulation over 3 days (**Fig. 2G**). Following this conditioning procedure, we observed that mice spent less time in the side in which they received photostimulation when allowed to explore both contexts (**Fig. 2H**). Interestingly, mice showed a significant increase in time spent in the center, transitional zone of the apparatus during the post-test as compared to both the pre-test and *vglut2*^VTA-LC::eYFP^ control mice (**Fig. 2I**). Additionally, *vglut2*^VTA-LC::ChR2^ mice showed a significant increase in the distance they traveled in the post-test session compared to pre-test (**Fig. 2J**). This increase in locomotor activity was also observed during photostimulation conditioning sessions as compared to non-stimulation conditioning sessions (**Fig. 2K**). During pre-test exploration, ChR2 mice showed no difference in center zone entries and greater velocity when locomoting in the center zone compared to either context (**Fig S2E&F**). However, this behavior is reversed during post-test exploration (more entries and no difference in velocity compared to either context) indicating a shift in preference. We also observed increased velocity between pre- and post-tests for ChR2 mice (**Fig. 2L**), but not between the conditioned and non-conditioned sides (**Fig. S2F**). The increase in activity during the post-test session also correlated with change in context preference behavior during the post-test (**Fig. S3G**). Further, location heatmaps show distinct conditioned side-avoidance and center zone preference not observed in during pre-test exploration (**Fig. 2M**). Together these data demonstrate that stimulation of *vglut2*^VTA-LC^ projections is sufficient to drive robust aversion behaviors.

**Figure 2.**
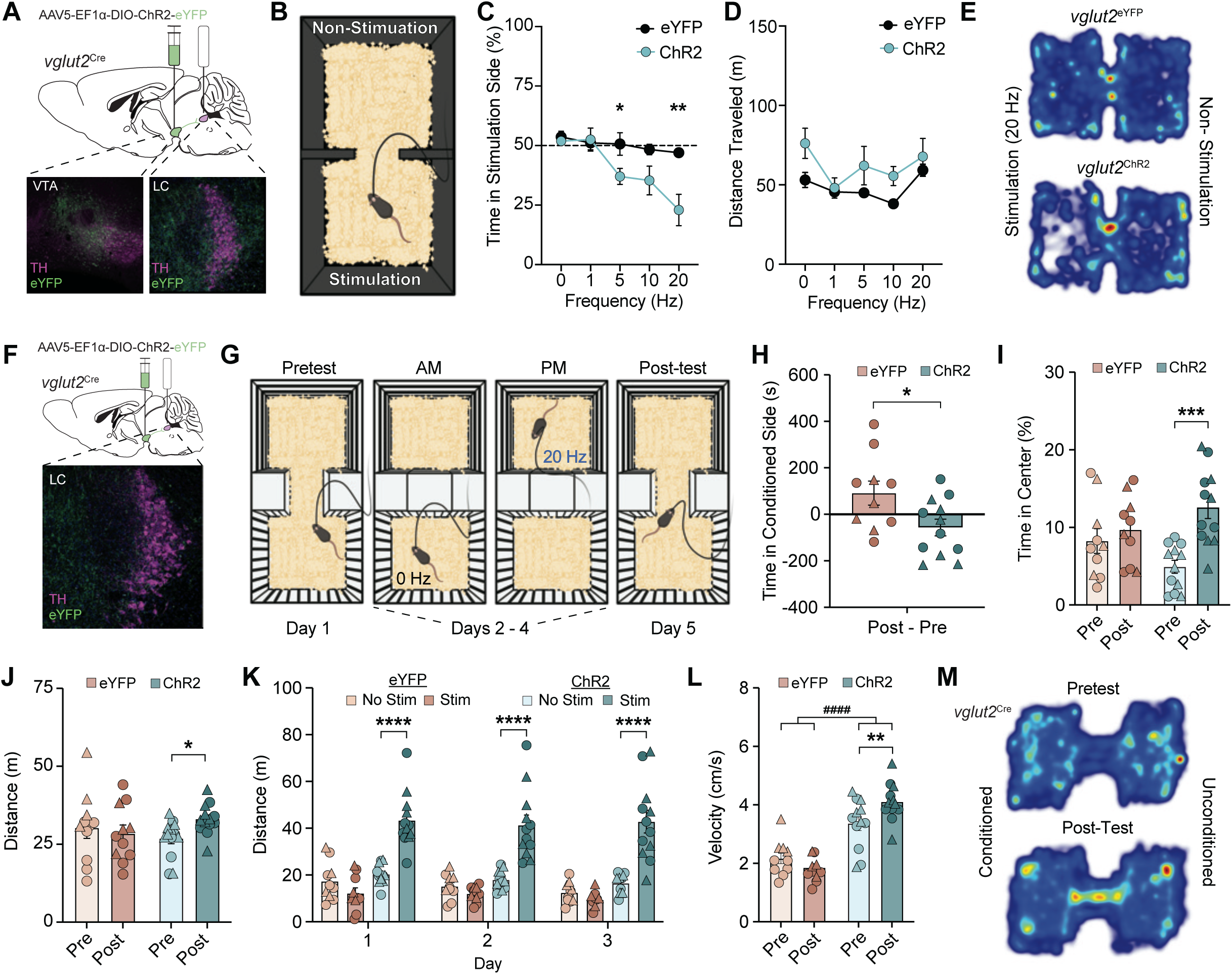
Photostimulation of *vglut2*^VTA–LC^ projections drives frequency-dependent real-time and conditioned place aversions. (**A**) Schematic of viral injection into the VTA and optic fiber implantation into the LC of *vglut2*^Cre^ mice (top) and representative coronal image of ChR2-eYFP (green) and tyrosine hydroxylase (TH, magenta) expression in the VTA (bottom left) and LC (bottom right). (**B**) Cartoon depicting optogenetic real-time place testing. (**C&D**) Increasing photostimulation frequency drives active avoidance in the real-time place test in ChR2-expressing *vglut2*^VTA–LC^ mice compared to eYFP controls (C; 2-way ANOVA: Stimulation x Group, F(4, 48) = 3.343, *p* = 0.0171; Uncorrected Fisher’s LSD: 5Hz, eYFP vs ChR2, p = 0.0383; 20 Hz, eYFP vs ChR2, p = 0.0099), but does not affect distance traveled (D; 2-way ANOVA: Stimulation x Group, F(4, 48) = 0.3808, *p* = 0.8212). (**E**) Representative heatmaps depicting location frequency in eYFP and ChR2-expressing *vglut2*^VTA–LC^ mice during 20 Hz stimulation. (**F**) Schematic of viral injection into the VTA and optic fiber implantation into the LC of *vglut2*^Cre^ mice (top) and representative coronal image of ChR2-eYFP (green) and tyrosine hydroxylase (TH, magenta) expression in the LC (bottom). (**G**) Cartoon depicting optogenetic conditioned place preference testing paradigm. (**H**) Repeated photostimulation (20 Hz) conditioning reduces time spent in conditioned side during post-test exploration ChR2-expressing *vglut2*^VTA–LC^ mice compared to eYFP controls (t(20)= 2.418, *p* = 0.025). (**I**) Repeated photostimulation (20 Hz) conditioning increases total time spent in center zone between chambers during post-test exploration in ChR2-expressing *vglut2*^VTA–LC^ mice (F(1, 20) = 5.548, *p* = 0.0288; Uncorrected Fischer’s LSD: ChR2, Pretest vs. Posttest, *p* = 0.0003). (**J**) Conditioning increases total distance traveled in both chambers during post-test exploration (2-way ANOVA: Group x Test Day, F(1, 22) = 4.66, *p* = 0.0419; Uncorrected Fisher’s LSD: ChR2, Pretest vs Posttest, *p* = 0.0225). (**K**) Photostimulation increases total distance traveled during conditioning days 1-3. (3-way ANOVA, Stimulation x Group: F(1,20) = 40.51, *p* < 0.0001); Tukey’s multiple comparison: ChR2 Day 1: No Stim vs. Stim *p* < 0.0001; ChR2 Day 2 : No Stim vs. Stim *p* < 0.0001; ChR2 Day 3: No Stim vs. Stim *p* < 0.0001). (**L**) Velocity is higher in ChR2 mice compared to control mice and after conditioning (2-way ANOVA, Group: F(1, 20) = 56.14, p <0.0001, Day x Group: F(1, 20) = 13.12, *p* = 0.0017, denoted as ####. Uncorrected Fischer’s LSD: ChR2, Pretest vs. Posttest, t(20) = 3.49, *p* = 0.0023; Control, Pretest vs. Posttest, t(20) = 1.7183.49, ns). (**M**) Heatmaps of average time spent in ChR2-expressing *vglut2*^VTA–LC^ mice during pre-test and post-test exploration. In all relevant panels, circles indicate male mice, triangles indicate female mice

### A modulatory role for synaptic integration of glutamatergic transmission from VTA to LC

The robust and rapid negative valence and conditioned aversion to neutral stimuli following photostimulation (**Fig. 2**) was somewhat surprising given the relatively small amplitude and rapid paired-pulse depression we observed in slice (**Fig. 1**). From this point we suspected that glutamatergic VTA-LC transmission might function to constructively summate with other excitatory synaptic inputs to the LC. To test this hypothesis, we combined non-selective electrical stimulation with photostimulation of *vglut2*^VTA-LC^ terminals in slice. To do so, we used a bipolar electrode to apply a train of electrical stimulation (10 pulses at 20 Hz) within the dendritic field of LC-NE neurons during a long-lasting *vglut2*^VTA-LC^ photostimulation at either 5 or 20 Hz (**Fig. 3A&B**). Generally, in acute slice preparation LC-NE neurons spontaneously fire around 1 Hz without external stimulation. The 5 second photostimulation alone (either 5 or 20 Hz) caused a mild increase of spontaneous firing frequency of LC-NE neurons. This enhancement started from a transient increase at the first second then reduced to a small but steady enhancement of firing rate (**Fig. 3C-E**). For current clamp recordings, we adjusted the intensity of electrical stimulation to have an approximate 10-50% probability of trial-to-trial success in inducing phasic activity of LC-NE neurons as in a prior study^59^. Interestingly, simultaneous electrical and optical stimulation significantly enhanced induction of phasic activity (**Fig. 3F-I**), but did not affect the spike number within each successfully evoked phasic burst (**Fig. 3J**). These results demonstrate integration between glutamatergic innervations from VTA to LC with general excitatory inputs to increase probability phasic burst activity. We next sought to test whether a similar phenomenon occurs *in vivo*.

**Figure 3.**
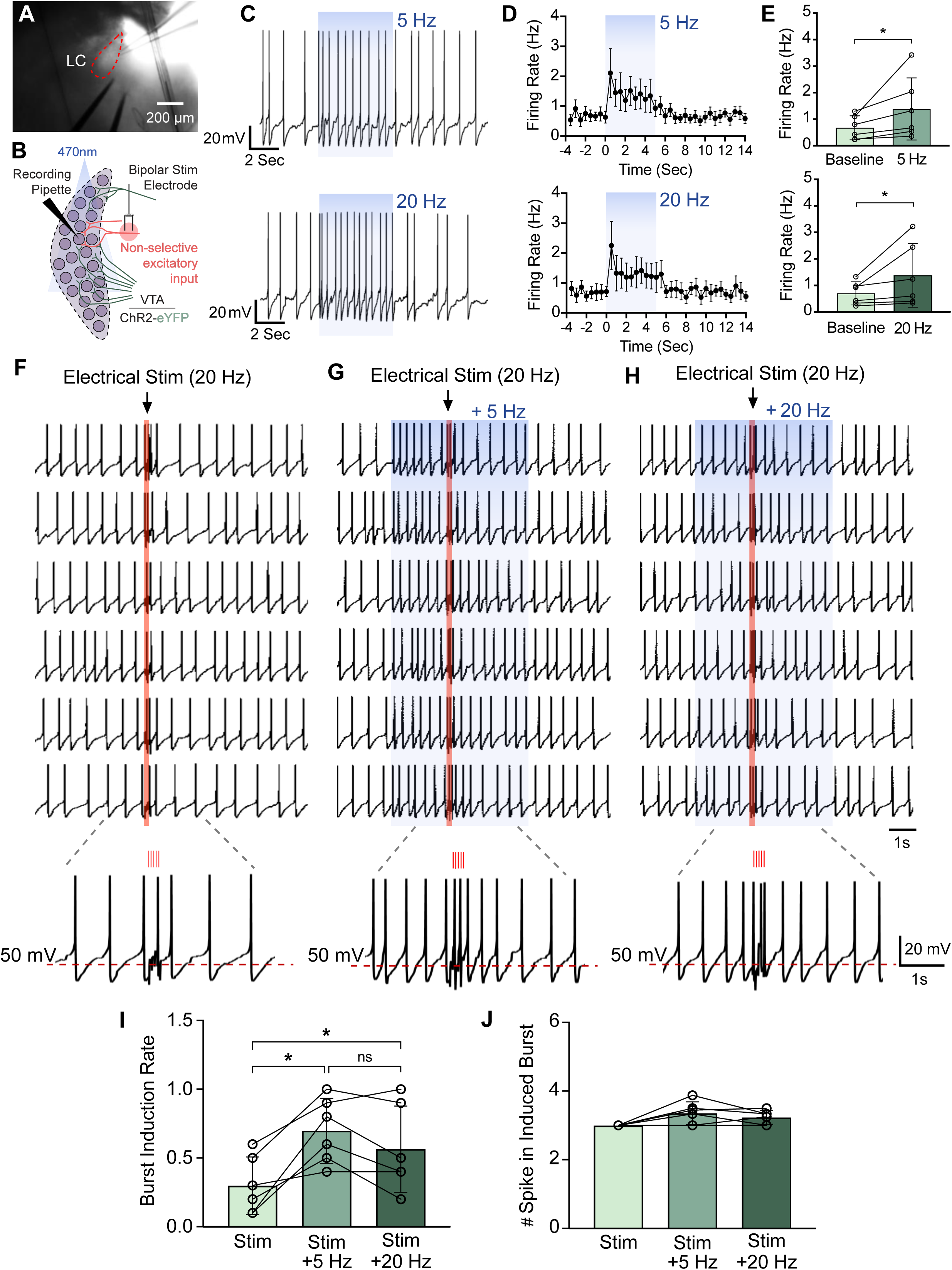
Coincident activation of *vglut2*^VTA–LC^ inputs integrates with other glutamatergic synaptic LC inputs to drive phasic bursts. (**A**) Low magnification DIC image demonstrating the placement of bipolar electrode. (**B**) Schematic illustrating experimental design. (**C-E**) Representative traces (C) and quantification (D) showing a steady augmentation of spontaneous firing rate upon the long-lasting photostimulation (5 s) at 5 Hz (top) and 20 Hz (bottom). Each stimulation paradigm significantly increased tonic firing (E; Wilcoxon matched-pairs signed rank test, 5Hz: W = 21, median of differences = 0.4154, p < 0.0312; 20Hz: W = 21, median of differences = 0.22, p < 0.0312). (**F-G**) Electrical stimulation (20 Hz) induces phasic activity in six consecutive sweeps without photostimulation (F), and with either 5 Hz(G) or 20 Hz (H) photostimulation. Electrical stimulation and photostimulation are denoted by red and blue rectangles, respectively. Peri-stimulus traces defined by dashed brackets demonstrate the induction of phasic activity. (**I**) Concurrent photostimulation significantly increases the burst induction rate (Repeated measures One-way ANOVA, F (1.824, 9.122) = 14.74, *p* = 0.0016, Tukey’s multiple comparisons: Stim vs Stim + 5 Hz, p = 0.0125; Stim vs Stim + 20 Hz, p = 0.0305). (**J)** Concurrent photostimulation did not alter spike number within each evoked burst (Friedman test, Friedman statistic = 5.444, p = 0.0741).

### *In vivo vglut2^VTA-LC^* photostimulation during action-associated punishment impairs task completion

To address whether *vglut2*^VTA-LC^ inputs facilitate negative affective behavior due to a selective amplification of excitatory synaptic inputs driven by noxious environmental stimuli, we employed a behavioral model that enables examination of discrete negative stimuli known to increase LC neuron activity and promote norepinephrine release^14,15,27,64–66^. Here we used the Punishment Risk Task (PRT), a task that uses unpredictable probabilistic punishment to delay, but not prevent, reward-seeking behavior. In particular, it measures the latency to pursue a reward while increased likelihood of action-associated punishment (foot shock) impacts this reward pursuit^67–69^. We sought to test whether *vglut2*^VTA-LC^ stimulation could impact behavioral responding to salient negative stimuli (foot shock) and exacerbate established anxiogenic responding (increased nosepoke latency). To better understand whether PRT engages VTA glutamate neurons, we injected a Cre-dependent fluorescent reporter virus (AAV5-eF1α-DIO-eYFP) into the VTA of *vglut2*^Cre^ mice (**Fig. 4A**) and trained them for PRT (**Fig. 4B-C**). Immediately following PRT testing, mice were euthanized, and their brains collected to examine cFos immunoreactivity, a proxy of neuronal activation. Histological examination of eYFP-expressing VTA neurons revealed increased percentage of co-expression of cFos immunoreactivity in eYFP+ VTA neurons in animals exposed to footshock during PRT as compared to animals given no foot shock during PRT (**Fig. 4D-G**). To determine if the *vglut2*^VTA-LC^ projection had any influence in this behavior, we injected AAV5-eF1a-DIO-ChR2-eYFP into the VTA of *vglut2*^Cre^ mice, implanted optic fibers above the LC and trained the mice in PRT (**Fig. 4H**). To mimic the electrical stimulation experiments in **Fig. 3**, we tested mice in multiple counterbalanced experimental conditions including varying shock amplitudes (0 mA, 0.05 mA and 0.1 mA), with or without concurrent photostimulation (0 or 20 Hz) (**Fig. 4I**). Here we found mice increased latency to respond to operant-associated cues when the probability of punishment increases (Block 1 vs Block 3), similar to previous studies^67–70^. We also determined that both 0.05 mA and 0.1 mA shock amplitudes were sufficient to drive increased nosepoke latency during increased probability of punishment (**Fig. 4J**). This latency was unaffected by 20 Hz photostimulation, except during the second block of 0.05 mA shock presentation. Here, shock-paired photostimulation increased nosepoke latency compared to 0.05 mA shock alone. Additionally, reward retrieval latency was also increased following increased probability of punishment (**Fig. 4K**) and was not affected by photostimulation. The average nosepoke latency described here is based on all committed nosepokes and did not account for animals that did not complete the task (45 trials). Following closer analysis, we observed that shock-paired photostimulation reduced the probability of task completion (**Fig. 4L**), specifically reducing the percentage of trials completed in the 3^rd^ Block (**Fig. 4M**). Analysis of average post-shock nosepoke latencies during the trial immediately following each shock revealed that combined 0.1 mA shock and 20 Hz photo-stimulation during the final shock significantly increased nosepoke latency (**Fig. S4A**). These data indicate that despite not significantly affecting the average nosepoke latency during Blocks 2 and 3, mice are significantly less likely to complete all 45 trials when *vglut2*^VTA-LC^ photostimulation is time-locked to shock presentation. Following PRT testing, we confirmed that these mice maintain the negative valence induced by *vglut2*^VTA-LC^ stimulation with real-time place testing. Here mice previously tested in PRT still actively avoided 10 Hz and 20 Hz photo-stimulation as observed in earlier experiments (**Fig. S4B**). However, we did not find any relationship between the magnitude of aversion (20 Hz) and the percentage of trials completed during combined stimulation (0.1 mA + 20 Hz) (**Fig. S4C**). Overall, the data presented here demonstrate *vglut2*^VTA-LC^ projections can enhance the aversive behaviors in response to noxious stimuli.

**Figure 4.**
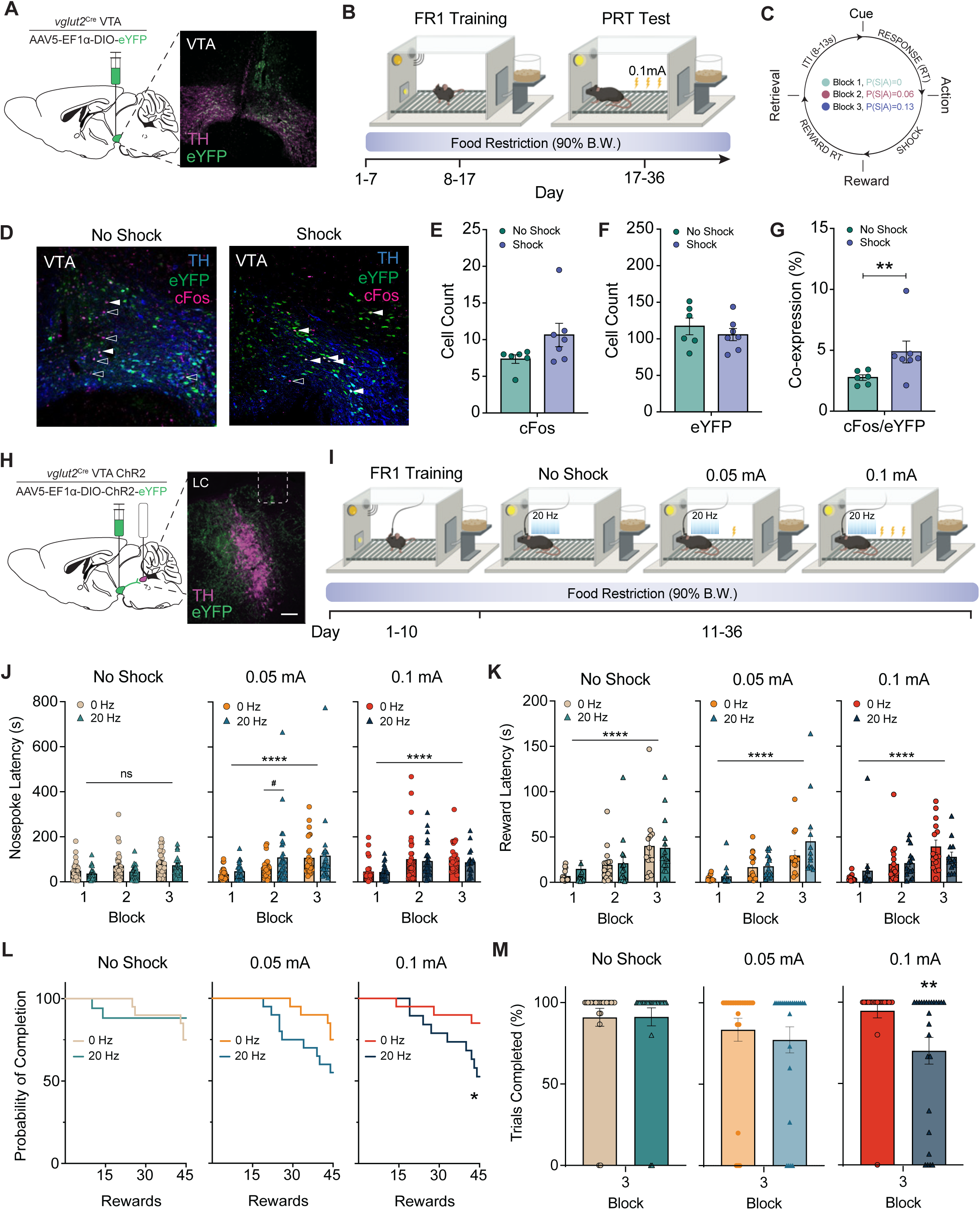
*vglut2*^VTA–LC^ activation exacerbates task failure in the Punishment Risk Test. (**A**) Schematic (left) of viral injection of ChR2-eYFP into the VTA and optic fiber implantation into the LC of *vglut2*^Cre^ mice and representative coronal image (right) of ChR2-eYFP expression (green) and tyrosine hydroxylase (TH; magenta) in the VTA. (**B**) Cartoon depicting punishment risk task (PRT) calendar. (**C**) Schematic depicting PRT task operant conditioning and shock presentation during trials with increasing probability of shock. (**D**) Representative histological images of VTA expression of cFos in *vglut2*^VTA::eYFP^ neurons in groups exposed to no shock (left) or shock (right). Images show eYFP (green), cFos (magenta), and TH (blue). (**E**) Total count of cells expressing cFos in the VTA is not changed by presence of shock during task performance (No Shock vs. Shock, Welch’s t test: t(1.936) = 7.535, *p* = 0.0911). (**F**) Total count of cells expressing eYFP in the VTA is not different between groups (No Shock vs. Shock, Welch’s t test: t(9.431) = 0.7986), *p* = 0.4442.). (**G**) Shock during task performance significantly increases percentage of *vglut2*^VTA^ neurons expressing cFos (Mann-Whitney test: U = 5, median (No Shock) = 2.787, n = 6; median (Shock) 4.348, n = 7), *p* = 0.0221). (**H**) Schematic of viral injection into the VTA and optic fiber implantation into the LC of *vglut2*^Cre^ mice (left) and representative coronal image of ChR2-eYFP (green) and tyrosine hydroxylase (TH, magenta) expression in the LC (right). (**I**) Cartoon and calendar depicting PRT photostimulation paradigm. (**J**) Increasing probability of shock increases nosepoke latency while photostimulation (20 Hz) or concurrent shock and photostimulation do not statistically alter nosepoke latency (Mixed effects model (REML): 0.05mA(Block), F(2, 62) = 14.12, *p* < 0.0001, Tukey’s multiple comparisons: Block 2, 0Hz vs 20 Hz *p* < 0.0399; 0.1mA(Block), F (1.363, 47.71) = 38.50, *p* < 0.0001. (**K**) Increasing probability of shock increases reward retrieval latency while photostimulation (20 Hz) or concurrent shock and photostimulation do not statistically change reward retrieval latency (Mixed effects model (REML): Block: F(1.503, 22.54) = 47.32, *p* < 0.0001. (**L**) Concurrent photostimulation significantly reduces the probability of PRT completion when mice are exposed to 0.1 mA shock (Log-rank (Mantel-Cox) test: χ² = 4.074, df = 1, *p* = 0.0351). (**M**) Photostimulation significantly reduces the percentage of trials completed in the 3^rd^ block when mice are exposed to 0.1 mA shock (Wilcoxon matched-pairs signed rank test, W = −45, *p* < 0.0039). In all relevant panels, circles indicate male mice, triangles indicate female mice

## Discussion

We examined the synaptic transmission and subsequent behavioral outputs driven by a relatively unknown and unstudied VTA-LC projection. Together our electrophysiological, optogenetic, and behavioral results support the premise that the glutamatergic VTA-LC projections form monosynaptic contacts onto LC-NE neurons. Further, we have demonstrated a unique role for this neural population in avoidance behaviors using the real-time place test, conditioned place test, and punishment risk task.

### Monosynaptic glutamatergic afferents to LC

We found that LC-projecting glutamatergic VTA neurons are monosynaptic in nature, expanding our understanding of previously described excitatory inputs to the LC. Other well-described afferents arise from the medial prefrontal cortex (mPFC), lateral hypothalamus (LH), ventrolateral periaqueductal gray (vlPAG), and rostral ventrolateral medulla (RVLM)^71–75^. There appear to be distinct anatomical patterns among direct glutamatergic projections to LC. Previous studies suggest innervation from mPFC and vlPAG targets the pericoeruleus region medial to LC in which LC-NE neurons organize their dendritic field^59,71,75^. LH and RVLM axons appear to innervate the LC core^71,72,75^. Interestingly, axonal arborizations from VTA show a broad distribution across dorsal pontine area without a clear target. The discrepancy in axonal distribution implies distinct postsynaptic LC-NE targets corresponding to the location of presynaptic components as well as possible synaptic integration when coincident transmission occurs. This notion is partially supported by the ultrastructure of LC dendrites showing close spatial proximity of presynaptic components from vlPAG to unidentified asymmetric synapses^74^. Surprisingly, the glutamatergic VTA-LC input has broader innervation to the majority of the dorsal pontine area but activation produces smaller amplitude oEPSCs in LC-NE neurons compared to prior examples in mPFC, vlPAG, LH and RVLM^71,72,75^, suggesting a possible modulatory role for VTA-LC inputs in synaptic integration. Here, in support of that notion, we show VTA-LC glutamatergic activation efficiently converts coincident subthreshold excitatory input into phasic LC bursts. Importantly, LC phasic discharge is thought to reorient attention in response to critical unexpected or salient environmental stimuli by adjusting the gain on discrete neural circuits. In this vein, the tight regulation of LC phasic activity ensures appropriate actions in response to different stimuli^76–78^. Taken together, the research suggests that the glutamatergic VTA-LC input enhances LC responsivity to weaker or neutral stimuli. Importantly, this idea was demonstrated using concurrent optical and electrical stimulation that enhanced burst generation in LC-NE neurons. Prior work showed simultaneous phasic transient optogenetic stimulation of LC-NE neurons coupled with low-intensity electrical paw stimulation enhanced somatosensory cortical neuron responses similar to high-intensity stimulation that induces innate phasic LC activitation^3^. Those findings support our argument of a dramatic change in behavioral outcomes from the promotion of LC phasic activity in response to less important stimuli.

### Possible contribution of co-released dopamine and neuropeptide transmission

Given the co-expression of *vglut*2 and TH in specific VTA subpopulations, co-release of dopamine from glutamatergic terminals from the VTA could also participate in synaptic input from this projection^44^(**Fig. S3**). However, in a separate study we found no pharmacological effects of applications of agonists targeting D1- and D2-like receptors in *ex vivo* recordings^79^, suggesting that dopaminergic transmission would not contribute to the synaptic currents measured here. Further examination is required to investigate the contribution of co-released transmitters from glutamatergic VTA-LC terminals in aversive behaviors. Despite intense efforts to understand the functional role of neuropeptidergic inputs to the LC using genetic or pharmacological tools targeting specific receptors^16–18,20,71,80–83^, little is known about the impact on behavioral outcomes arising from co-incident neuropeptide transmission to the LC^44,71,84,85^. To that end, a recent elegant study demonstrated that distinct glutamatergic inputs to the LC are differentially presynaptically regulated by bath application of neuropeptides known to act in the LC^71^. Naturally, multiple LC innervations could be concurrently activated by salient or noxious stimuli and converge to promote aversive behavior. It is possible that glutamatergic VTA-LC transmission may help shape the impact of these coincident neuropeptidergic inputs. Further studies will be required to understand the interaction between these different LC afferents during distinct behaviors.

### VTA glutamate neurons drive negative affect behavior and lead to failure in punishment-associated actions

Given the LC-NE system’s important role in modulating negative affective states^4–11,16,19,23,24,26,86–88^, we used *in vivo* optogenetics to determine the effect of VTA afferent stimulation in the real-time place and conditioned place preference tests. Here we found that animals actively avoided real-time stimulation and spent less time in stimulation-conditioned contexts indicating negative affective behavior. These data parallel previously described results of optogenetic and chemogenetic stimulation of LC-NE neurocircuitry^16,19,23,26,86–88^. However, these strategies inherently raise the possibility of co-activating bypassing axons and driving synaptic release in collaterals via antidromic action potentials^89^. Additionally, photostimulation of VTA-LC glutamatergic afferent could also recruit local circuitry that could further the synaptic input^59,90^. Another noteworthy outcome in the CPP testing was that mice also spent more time in the center passageway between contexts, where no specific contextual cues were present. We suspect that conditioning may have promoted a negative affective association to the entire testing environment and the shift to time spent in the center may be a behavioral adaptation that generalized aversion to either context. The functional diversity of LC-NE efferents likely has differential effects on fear conditioning and fear generalization dependent upon the target site^24^. Taking these findings and our electrophysiological results, we used the Punishment Risk Task (PRT) to test whether this excitatory input had any influence on risk associated hesitancy to pursue a food reward. This model enables assessment of a complex negative affective state that develops when goal-directed actions have a low probability of an aversive outcome. Importantly, this task involves the sensitivity to a discrete noxious stimulus (foot shock) known to increase LC-NE activity and related noradrenergic action^14,15,27,64–66^. Here, the presence of this noxious stimulus during positive reinforcement disrupts typical low latency behavioral responding by delaying these reward-associated actions without abruptly preventing them. We previously characterized this behavior as a measure of “hesitancy” or “caution” during this reward seeking behavior^70^. A recent study also found that mice delayed their action for a signaled reward when a new, unsignaled rule that punished the action was added^91^. Further, anxiolytic pharmacological interventions are also effective in eliminating any behavioral changes driven by probabilistic punishment^67,68,70^. For this study, PRT was an ideal behavioral task to test the hypothesis that arose from our electrophysiology results in **Fig. 3**. Namely, here we could vary the shock amplitude in an analagous manner to the coincident electrical stimulation. Animals were trained in the PRT task and then tested with different amplitudes of shock and coinciding optogenetic stimulation. When mice received *vglut2*^VTA-LC^ photostimulation (20 Hz) that coincided with footshock, we found an increased probability of animals not completing the task (45 trials). Specifically, when photostimulation is paired to 0.1mA footshock, mice completed fewer trials compared to 0.1 mA footshock alone. This trend was also observed in our previous study data that showed higher amplitude footshock (0.3 mA) resulted in more animals not completing the task as compared to 0.1 mA^70^. Similarity of these outcomes may be due to similar increases in LC activity following footshock induction. Indeed, other studies have demonstrated that incremental increases of footshock intensity drive incremental increases in LC activity^3,14,15,25,64–66^ and important recent accumulating evidence across species has suggested the LC functions as a global model failure system leading to altered behavior when expectations about the environment are robustly refuted^92–94^. In line with this concept, our findings suggest that *vglut2*^VTA-LC^ photostimulation may increase LC signal integration in a way that amplifies the salience of mild footshocks to reduce task completion.

Altogether, we identify a monosynaptic glutamatergic VTA projection to the LC that is well-positioned to modulate other excitatory input to the LC. Activation of this projection enhances avoidance behaviors that guide actions away from risk-associated outcomes. Understanding how this distinct glutamatergic VTA efferent regulates LC-mediated negative affect and subsequent behaviors will important insights into the heterogeneity and diverse behavioral functions of both VTA glutamate and LC-NE neurons. Ultimately, these insights could be essential to understanding how diverse neurocircuitry modulate LC activity to coordinate competitive motivational states dysregulated in various physiological or pathological conditions including anxiety, depression, substance use disorder, and pain.

## Methods

### Subjects

Male and female C57BL/6J (RRID:IMSR_JAX:000664) and vGlut2^Cre^ (RRID:IMSR_JAX:016963) 3-6 month old mice were bred locally. Mice were group-housed, given access to food pellets and water ad libitum, and maintained on a 12 hr:12 hr light:dark cycle (lights on at 7:00 AM). Mice were transferred to the experimental facility and allowed 1 week of habituation. Animals were gently handled and weighed weekly.

### Stereotaxic surgery

Following acclimation to the holding facility for at least seven days, the mice were anesthetized in an induction chamber (3% isoflurane) and placed into a stereotaxic frame (Kopf Instruments, Model 1900) where they were maintained at 1-2% isoflurane. For electrophysiological experiments and behavioral experiments, mice were then injected unilaterally (250 nL, Neuros Syringe, Model 7000.5, Hamilton) at a rate of 50nL/min in the VTA at stereotaxic coordinates: AP −3.1mm, ML 0.5mm, DV −4.5mm). Viruses used include AAV5-EF1a-DIO-hChR2-(H134R)-eYFP, AAV5-EF1a-DIO-eYFP, AAV5-CaMKII-eYFP and AAV5-CaMKII-ChR2-eYFP (The Hope Center Viral Core, Washington University, St. Louis, MO). Optic fiber implants were directed to the LC at stereotaxic coordinates: AP −5.4, ML 0.8, DV −3.25. Mice were allowed to recover for five weeks prior to electrophysiological or behavioral examination, permitting optimal expression of the virus. For optogenetic experiments, an additional intracranial optic fiber implant was directed above the LC (AP −5.4, ML: 0.8, DV −3.25) following injection. Implants were secured using Metabond Quick Adhesive Cement and gel superglue.

### Brain slice preparation and electrophysiology

Adult *vglut2*^VTA-LC::ChR2^ mice were anesthetized with a cocktail of ketamine, xylazine & acepromazine then perfused with slicing-aCSF consisting of (in mM) 92 N-methyl-d-glucose (NMDG), 2.5 KCl, 1.25 NaH_2_PO_4_, 10 MgSO_4_, 20 HEPES, 30 NaHCO_3_, 25 glucose, 0.5 CaCl_2_, 5 sodium ascorbate and 3 sodium pyruvate, oxygenated with 95% O2 and 5% CO2, pH 7.3–7.4, and osmolality adjusted to 315–320 mOsm with sucrose. Brains were block mounted with 2% agarose made in slicing-aCSF and coronal brainstem sections in 350 μm thickness were cut through a vibratome (VF310-0Z, Precisionary Instruments, MA, USA). Slices were incubated in warm (32°C) slicing-aCSF for 30 minutes then transferred to holding-aCSF that consisted of (in mM) 92 NaCl, 2.5 KCl, 1.25 NaH_2_PO4, 30 NaHCO_3_, 20 HEPES, 25 glucose, 2 MgSO_4_, 2 CaCl_2_, 5 sodium ascorbate and 3 sodium pyruvate, oxygenated with 95% O2 and 5% CO2, pH 7.3–7.4, and osmolality adjusted to 310–315 mOsm. Under the visual guidance from a digital CMOS camera (ORCA-Flash4.0LT, Hamamatsu Photonics, Shizuoka, Japan) and a 40x water immersion objective lens (LUMPLFLN-40xW, Olympus, Tokyo, Japan), whole cell recordings were made in a recording chamber mounted on an upright microscope (BX51WI, Olympus Optical Co., Ltd, Tokyo, Japan) and perfused with warm (29–31°C) recording-aCSF containing (in mM) 124 NaCl, 2.5 KCl, 1.25 NaH_2_PO_4_, 24 NaHCO_3_, 5 HEPES, 12.5 glucose, 2 MgCl_2_, 2 CaCl_2_, oxygenated with 95% O2 and 5% CO2, pH 7.3–7.4, and osmolality adjusted to 315–320 mOsm with sucrose. Glass pipettes pulled by borosilicate glass capillary (GC150F-10, Warner Instruments, Hamden, CT, USA) with a resistance around 10 MΩ were filled with potassium gluconate-based intra-pipette solution consisting of (in mM) 120 potassium gluconate, 5 NaCl, 10 HEPES, 1.1 EGTA, 15 Phosphocreatine, 2 ATP and 0.3 GTP, pH 7.2–7.3, and osmolality 300 mOsm. Recordings under voltage clamp mode were held at −70mV unless specified and be discarded if the serial resistance varied greater than 20% of the baseline value that should be less than 20 MΩ. Current clamp recordings were discarded when the membrane potential (Vm) was over −40 mV or the APs were not able to overshoot at 0 mV. All of data were collected by Multiclamp 700B amplifier (Molecular Devices, San Jose, CA, USA) under a low-pass filtered at 2 kHz and digitized at 10k Hz by Axon Digidata 1550B interface (Molecular Devices, CA, USA) coupled with Clampex software (Molecular Devices, CA, USA). In optogenetic experiments, 2 ms light pulses at 10 mW (unless specified), generated by LED light source (Cyclops LED Driver, OpenEphys, GA, USA) and delivered through epifluorescence equipment mounted on the microscope every 10 or 60 seconds for long-lasting photostimulation at either 5 or 20 Hz. For electrical stimulation, 5 electrical pulses at 20 Hz (duration: 500 μs; intensity: 20–80 μA) every 40 seconds were generated by a stimulus isolator (A385, World Precision Instruments, FL, USA) and delivered through a bipolar tungsten electrode (FHC, Inc., Bowdoin, ME, USA), which was placed in the medial dendritic field of LC. Both electro- and photo-stimulations were commanded by Axon Digidata 1550B interface. The criteria of detection of evoked phasic activity in LC-NE cells was modified from prior work^59^. Namely, an evoked phasic activity includes at least 3 consecutive spikes with instantaneous frequency over 5 Hz, which should exceed the value of mean + 2 SD of firing rate in baseline epochs based on the classical observation of phasic activity in *in vivo* studies^14,95^. Pharmacological applications were applied with recording-aCSF through perfusion system. All electrophysiological data were gathered and further analyzed by Clampex software, Matlab (MathWorks, MA, USA) and GraphPad Prism (GraphPad Software, MA, USA).

### Immunohistochemistry

Mice were intracardially perfused with 4% paraformaldehyde, and then brains were sectioned (30 μm) and placed in 0.1 M PB until immunohistochemistry. Free-floating sections were washed in 0.1 M PBS for 3 × 10 minutes intervals. Sections were then placed in blocking buffer (0.5% Triton X-100 and 5% natural goat serum in 0.1 M PBS) for 1 hr at room temperature. After blocking buffer, sections were placed in primary antibody (1:1000 chicken anti-TH, Aves Labs; 1:1000 chicken anti-GFP (GP1010), Aves Labs; 1:500 mouse anti-TH (MAB318), MilliporeSigma; 1:1000 rabbit anti-mCherry (600-401-P16), Rockland) in 10% blocking buffer overnight at room temperature. After 3 × 10 minutes 0.1 M PBS washes, sections were incubated in secondary antibody (AlexaFluor 488 conjugated goat anti-chicken IgG, AlexaFluor 594 conjugated goat anti-rabbit IgG, AlexaFluor 633 conjugated goat anti-mouse IgG, AlexaFluor 594 or AlexaFluor 633 goat anti-chicken IgG, Life Technologies) for 2 hours at room temperature, followed by subsequent washes (3 × 10 minutes in 0.1 M PBS, then 3 × 10 minutes 0.1 M PB washes). After immunostaining, sections were mounted and coverslipped with Vectashield HardSet mounting medium with DAPI (Vector Laboratories) and imaged on a Leica TCS SPE confocal microscope.

## Behavior

### Real-Time Place Testing

We used custom-made unbiased, balanced two-compartment conditioning apparatus (52.5 x 25.5 x 25.5 cm) as described previously^16,23,96,97^. Mice were allowed to freely roam the entire apparatus for 30 min. Entry into one compartment triggered constant photo-stimulation at either 1 Hz, 5 Hz, 10 Hz, or 20 Hz (473 nm, 10 ms pulse width, ∼10 mW light power) while the animal remained in the light paired chamber. Entry into the other chamber ended the photo-stimulation. The side paired with photo-stimulation was counterbalanced across mice. Time spent in each chamber and total distance traveled for the entire 30-minute trial was measured using EthovisionXT 13 (Noldus Information Technologies, Leesburg, VA).

### Conditioned Place Preference

We used a modified three-chamber CPP apparatus consisting of two square boxes (27 cm × 27 cm) that served as the conditioning chambers separated by a small center area that served as the passageway (5 cm wide × 8 cm long) between boxes. Boxes had 2.5 cm black-and-white vertical stripes or horizontal stripes and floors were covered with 500 ml of bedding on each side. The Plexiglas floor of the apparatus was covered with corncob bedding. Mice were transported to the CPP behavior testing room and handled once per day for at least 7 d before behavioral testing. Mice were then conditioned using a counterbalanced CPP. On day one, mice were allowed to explore all three regions of the box 30 minutes. Mice were paired in a counterbalanced fashion. Conditioning occurred over the following 3 days in which mice received no stimulation in the morning while confined to one side of the CPP box for 30 min. In the afternoon, at least 4 h after the morning conditioning session, mice were confined to the opposite side for 30 min and received one 20 Hz pulse of photo-stimulation every 30 seconds. The following day, mice were allowed to explore the entire apparatus and following the same procedure as the pretest. Preference scores were calculated by subtracting time spent in the stimulation-paired side during the pretest from time spent in the stimulation-paired side during the posttest.

### Open Field Test

OFT testing was performed in a sound attenuated behavior testing room as previously described. Lighting was stabilized at ∼12 lux. Animals were allowed to habituate to the room for 1 hour prior to testing. Each animal was individually placed into a 2500 cm^2^ enclosure for 20 minutes. The center of the apparatus was defined as a square equal to 25% of the total area. Activity was video recorded via a Google Pixel 3 XL and videos analyzed using Ethovision XT 13 (Noldus Information Technology, Wageningen, The Netherlands). Distance traveled and time spent in the center and periphery zones of the apparatus were determined and averaged for each animal.

### Elevated Plus Maze

EPM testing was performed within a sound attenuated behavior testing room. Lighting was stabilized at ∼12 lux. Animals were allowed to habituate to the room for 1 hour prior to testing. Each animal was individually placed into a plus-shaped platform for 15 minutes. The apparatus is comprised of two open arms (25 x 5 x 0.5 cm) across from each other and perpendicular to two closed arms (25 x 5 x 16 cm) with a center platform (5 x 5 x 0.5 cm) all 50 cm above the floor. Activity was video recorded via a Google Pixel 3 XL and videos analyzed using Ethovision XT 13 (Noldus Information Technology, Wageningen, The Netherlands). Distance traveled and time spent in the open and closed arms was determined and averaged for each animal.

### Punishment Risk Task

The mouse Punishment Risk Task (PRT) was performed within sound-attenuated boxes (Med Associates Inc., Fairfax, VT) as described^70^. Prior to PRT training and testing, mice were handled and weighed daily and placed under a restricted diet to maintain 90% of free feeding bodyweight. Mice were placed into their respective cages and given equal weight pellets (2.5g for males, 2.0g for females) and observed to confirm consumption. After one week, animals were trained for PRT. Mice had free access to water. Mice were maintained on a 12:12-hr light/dark cycle (lights on at 7:00 AM). Animals were weighed daily to confirm and maintain food restricted weight throughout the experiment. Mice were trained to nosepoke on an FR1 schedule in response to a combined auditory (1s, 1000 Hz) and light cue inside the associated cue port to receive a sweetened pellet (Dustless Precisio– Pellet - F0071, Bio-Serv, Flemington, NJ). The light cue remained on until nosepoke completion. Completion of nosepoke entry was determined by disruption of an infrared activity monitor located inside the nosepoke port. Immediately following completion of the nosepoke, the port light turned off and a house light illuminated the chamber, and a sweetened food pellet was dispensed into the food hopper. Once the mouse entered the food hopper to retrieve the pellet, the house light turned off and a randomized intertrial interval of 8-13 seconds was initiated, followed by the next trial and cue presentation. The nosepoke port and food hopper were located on opposite walls of the operant chamber. The fixed ratio remained at one throughout all training and test sessions. Specifically, each mouse received one sweetened pellet after every completed nosepoke entry. Each mouse completed daily training sessions of 45 trials (60 minutes session duration, 10 days total) prior to the testing session. After training sessions, mice underwent a no-shock control test session, throughout which they performed 45 trials of nosepokes divided into three blocks separated by 2 minutes of darkness without any probability of punishment. Next, mice were randomly assigned to treatment groups to assess action-associated punishment behavioral responses. Mice received one of three footshock amplitudes (0mA, 0.05mA, or 0.1mA, 300 ms) and either 20 Hz photo-stimulation or no photo-stimulation, resulting in 6 possible treatments per mouse. Each block consisted of 15 trials with the associated contingency value assigned to each block in ascending order of probability. That is, in Block 1 there was a 0% chance of foot shock, in Block 2 there was a 6.66% chance of foot shock, and in Block 3 there was a 13.33% chance of foot shock. Each block was separated by 2 minutes of darkness to help discern a change in block. In Block 2, mice were randomly assigned to received one shock upon nosepoke number 2, 3, or 4. In Block 3 mice were randomly assigned to received one foot shock upon nosepoke number 2, 3, or 4 then a second foot shock upon nosepoke number 6, 7, 8, or 9. This pseudo-randomization of the shock presentation follows the shock parameters used in Park and Moghaddam (2017). As previously described^70^, we used a maximum cutoff of 60 minutes for each block for all training and test sessions. Sessions that were terminated at this cutoff were excluded. The number of mice and session that were terminated was recorded. We did not observe habituation to the foot shock as evidenced by lack of behavioral changes across sessions. Animals that did not reach training criteria or complete the first trial block within 60 minutes were excluded. Throughout all complete sessions, nosepoke latency and reward latency was recorded. Nosepoke latency refers to time between auditory/light cue onset and completed nosepoke and reward latency refers to time between reward delivery and reward retrieval. All procedures were approved by the Institutional Animal Care and Use Committee of Washington University, conformed to US National Institutes of Health guidelines, and the ARRIVE guidelines (Animal Research: Reporting of In Vivo Experiments) were followed as closely as possible. All procedures were approved by the Animal Care and Use Committee of Washington University and conformed to US National Institutes of Health guidelines.

#### Statistical analyses

For statistics of all electrophysiological results, the normality of data was first tested through Shapiro–Wilk test. Student’s t-test or paired t-test was used when both groups reach normal distributions; otherwise, the Mann–Whitney test or paired-Wilcoxon-sign rank test was used. Repeated measures one-way ANOVA followed by post-hoc Tukey’s test were used for comparisons among multiple groups (**Fig. 1E, 3G**). Statistical analyses were performed in GraphPad Prism 10.0. All data are represented as mean ± SD except Fig. 3D showing mean ± SEM.

## Acknowledgements

We thank the other members of the Al-Hasani and McCall labs, particularly Jenny R. Kim and Rui-Ni Wu for helpful feedback on this project. This work was financially supported by the Brain & Behavior Research Foundation (NARSAD YI – 28565, J.G.M.), the National Institutes of Health (R21DA060414; R01NS117899; R01NS135401, J.G.M.), and the McDonnell Center for Systems Neuroscience (K.P.). We would like to acknowledge biorender.com for figure cartoons, the Washington University School of Medicine Hope Center for Neurological Disorders viral vector core, and the Osage Nation, Missouria, Illinois Confederacy and many other tribes as the ancestral, traditional, and contemporary custodians of the land where this work was conducted.

## Author contributions

K.E.P., C-C.K., A.R.B., and J.G.M. conceived the project and designed the detailed experimental protocols. K.E.P., C-C.K., A.R.B., A.P.P., V.D., and S.C.H performed the mouse experiments. K.E.P., C-C.K., A.R.B., and J.G.M performed the investigation and analyzed the data. K.E.P., C-C.K., A.R.B., and J.G.M wrote the paper. K.E.P., C-C.K., and J.G.M. edited the paper. K.E.P., and J.G.M. acquired funding. J.G.M. provided research supervision. J.G.M. led overall project administration. All authors discussed the results and contributed to revision of the manuscript.

## Conflict of Interest

The authors declare no conflicts of interest.

---

**Supplementary Figure 1.**
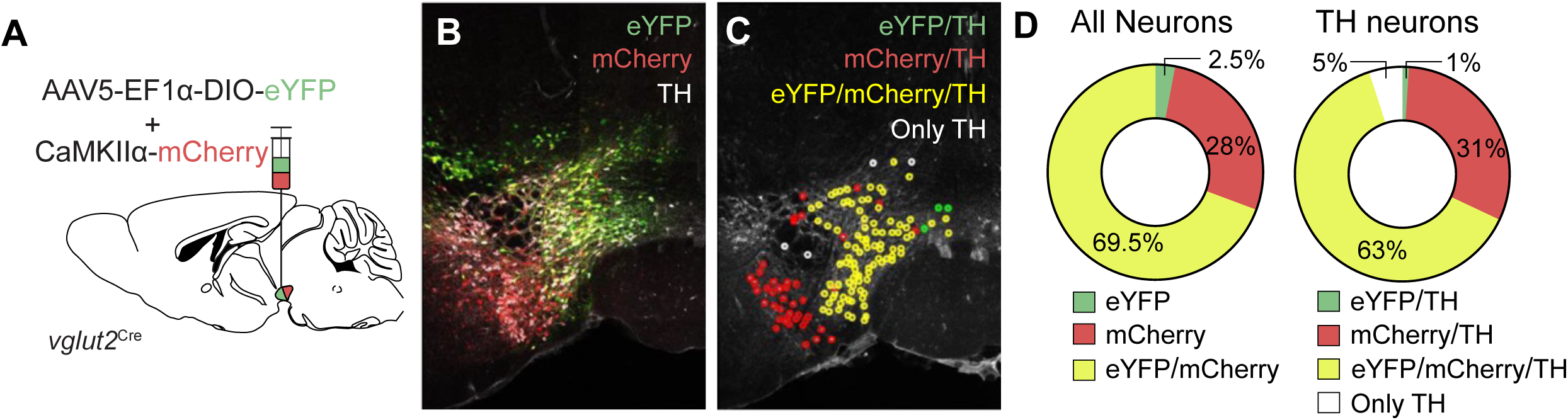
Distribution of *vglut2* and CaMKIIα-expressing neurons in the VTA. (**A**) Schematic of viral injection into the VTA of *vglut2*^Cre^ mice. (**B&C**) Confocal images showing the distribution of *vglut2*^Cre^-driven eYFP (green) expression, CaMKIIα driven mCherry expression (red), and TH-immunoreactive signals (white). Markers (C) indicate counted cells, and color indicates cellular identity. (D) Summarized results of cellular identity among *vglut2*^Cre^- and CaMKIIα-driven expression (left) and their colocalization with TH-immunoreactive signals (right).

**Supplementary Figure 2:**
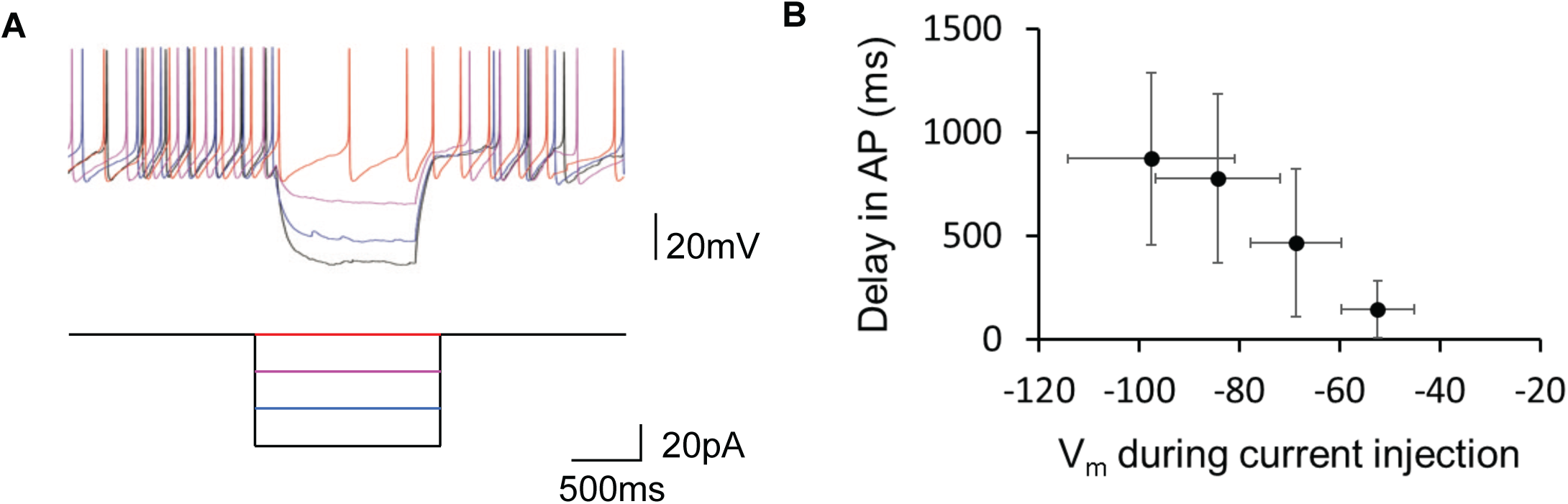
Electrophysiological identification of LC-NE neurons from delayed spiking after hyperpolarization. (**A**) Representative current-clamp recording from an LC-NE cell (top) in response to hyperpolarizing current injections (bottom). (**B**) Summarized results demonstrate the dependence on hyperpolarized membrane potential for delayed AP firing.

**Supplementary Figure 3.**
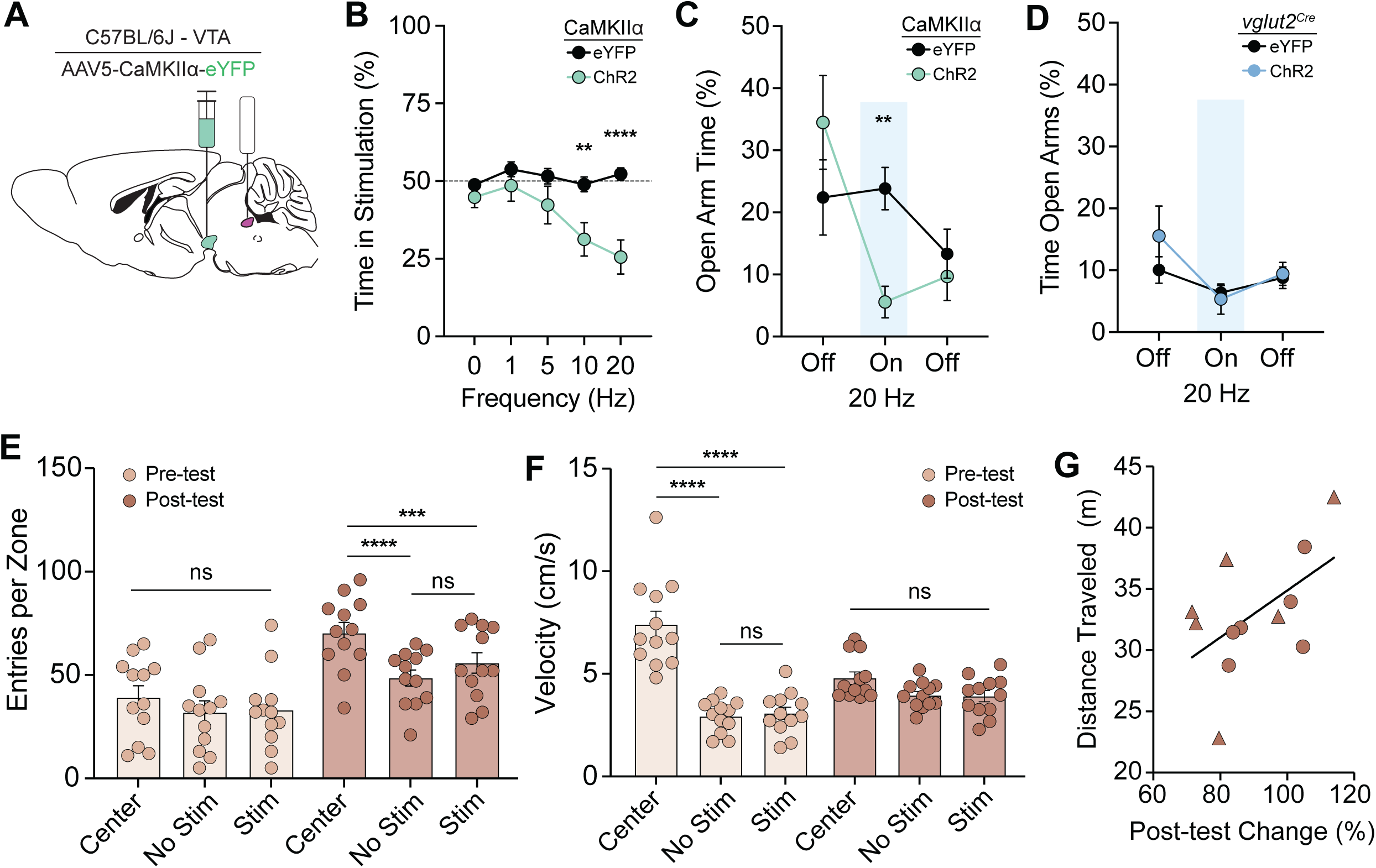
Photostimulation of CaMKIIα^VTA–LC^ projections drives frequency-dependent real-time place aversion and avoidance behavior. (**A**) Schematic depicting AAV5-CaMKIIα-eYFP injection into the VTA optic fiber implantation into the LC of C57BL/6J mice. (**B**) Increasing frequency of photostimulation drives active avoidance in real-time place test compared to eYFP controls (2-way ANOVA: Stimulation x Group, F (4, 56) = 4.083, *p* = 0.0057; Tukey’s Multiple comparisons test: 10Hz, eYFP vs ChR2, *p* = 0.0022; 20 Hz, eYFP vs ChR2, *p* < 0.0001). (**C**) Photostimulation (20 Hz) reduces time spent in the open arm of the elevated plus maze for CaMKIIα-ChR2-expressing mice during stimulation (Paired t test: ChR2(Off vs On), t(7)=3.69, *p* = 0.0078. (**D**) Photo-stimulation (20 Hz) has no effect on open arm exploration in ChR2-expressing *vglut2*^VTA–LC^ mice compared to *vglut2*^VTA–LC^ eYFP controls (2-way repeated measures ANOVA: F (2, 22) = 0.1735, *p* = 0.8419. (**E**) *vglut2*^VTA–LC::ChR2^ mice have greater average number of Center Zone entries during the CPP post-test (2-way ANOVA: Day x Zone, F (2, 44) = 4.165, *p* = 0.0221; Tukey’s multiple comparisons test, Center vs. NON-STIM, *p* < 0.0001, Center vs. STIM, *p* = 0.0005). (**F**) Center Zone is higher compared to the other zones during the CPP pre-test (2-way ANOVA: Day x Zone, F (2, 22) = 29.24, *p* < 0.0001; Center vs. NON-STIM, *p* < 0.0001, Center vs. STIM, *p* < 0.0001). (**G**) Distance traveled during the post-test has a postive relationship with the change in post-test in *vglut2*^VTA–LC::ChR2^ (Simple linear regression: F(1,10) = 3.976, *p* = 0.0741).

**Supplementary Figure 4.**
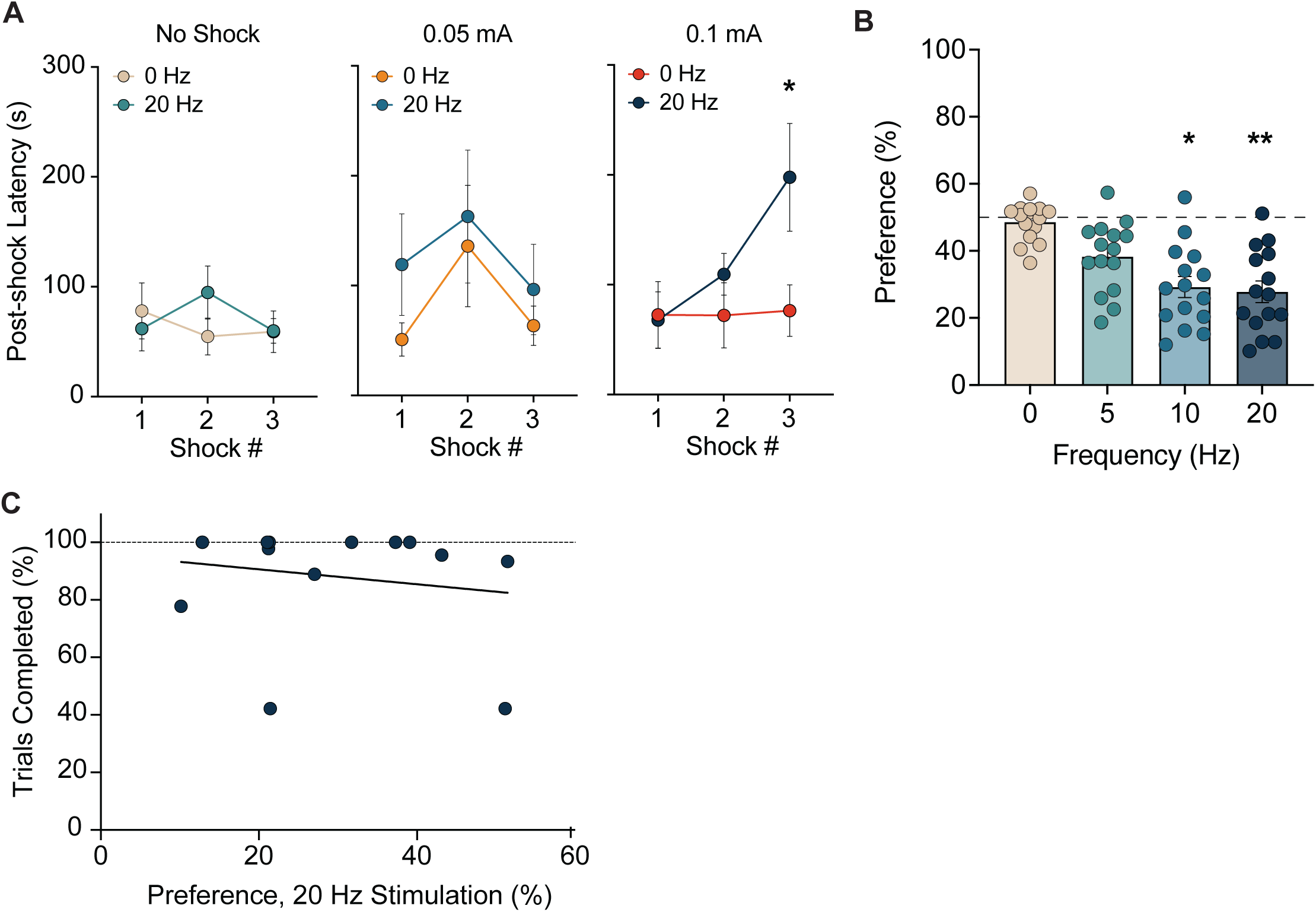
Punishment risk task completion does not correlate with real-time place preference. (**A**) Nosepoke latency for the trial immediately following shock, photostimulation, or simultaneous shock and photostimulation shows that concident shock and photostimulation enhances post-shock latency (Mixed-effects model (REML): Stimulation: F(1, 15) = 6.498, *p* = 0.0222; Shock Number: F(1.793, 26.89) = 5.716, *p* = 0.0104; Tukey’s multiple comparisons test, 0.1mA + 0Hz vs. 0.1mA + 20Hz, *p* = 0.0343). (**B**) Increasing frequency of photo-stimulation drives active avoidance in real-time place test in PRT-trained mice (One-way ANOVA: F(3,56) = 7.147, *p* = 0.0004). (**C**) Correlation of RTPT aversion (20Hz) and percentage of trials completed during combined stimulation (0.1 mA + 20 Hz) (Simple linear regression: F(1,12) = .3793, *p* = 0.54995).

## Notes

### Competing Interest Statement

The authors have declared no competing interest.

### Summary of Updates

We have updated the text including the abstract, introduction, discussion, and figure legends.

